# Steric attenuation of membrane-associated dimerization-induced fluorescence for cell-free biosensing

**DOI:** 10.64898/2026.10.19.760250

**Authors:** Parsa Parvizian, Hossein Moghimianavval, Allen P. Liu

## Abstract

We present a membrane-associated, cell-free biosensing reporter based on rapalog-induced split sfCherry reconstitution. Rapalog increased fluorescence in solution, lipid vesicles, and supported lipid bilayers. Antibodies binding to epitope tags near the dimerization interface reduced the fluorescence signal, consistent with steric interference with reporter assembly.

---

Biosensors are important tools in clinical diagnostics, environmental monitoring, and biosecurity^1^. Whole-cell biosensors are attractive due to their high specificity and cost-effectiveness, but their application has been limited by challenges in maintaining cell viability and stability^2^. To overcome the limitations posed by cellular viability, cell-free expression (CFE) systems have emerged as a powerful alternative, offering a highly programmable environment for molecular engineering^3^. When integrated with synthetic lipid compartments, CFE systems can move beyond simple protein expression to construct non-living cell-like systems capable of programmed environmental sensing and communication^4^.

Although cell-free biosensors offer programmable and portable detection capabilities^5,6^, conventional antibody-based assays such as ELISA remain widely used for specific biomarker detection^7^. ELISA workflows usually require sequential incubation, washing, and endpoint detection steps, making them relatively time-consuming assays^8^. Homogeneous solution-based cell-free sensors can simplify operation by allowing direct access to the analyte. Cell-free systems can also be combined with lipid membranes to construct membrane-associated sensing systems, extending their capabilities beyond soluble reporters^9^. However, compartmentalizing CFE systems inside phospholipid vesicles introduces mass-transfer constraints. This is particularly relevant to biosensors designed to detect large ligands, as large soluble proteins generally do not passively cross phospholipid bilayers^10^.

These trade-offs motivate the development of synthetic membrane biosensors that combine the programmability of cell-free systems with the spatial organization of a membrane interface. Living-cell reporters provide native membrane organization, but their outputs can be influenced by cell viability, endogenous signaling, membrane transport, and expression variability. Vesicle-based systems such as giant unilamellar vesicles (GUVs)^11^ partially address these limitations by providing biomimetic membrane compartments; however, as mentioned before, encapsulated reporters can remain inaccessible to membrane-impermeable analytes, and their sensitivity to mechanical perturbations can complicate handling^12^. We therefore sought to build a vesicle-inspired reporter platform in which cell-free-expressed sensing components are displayed on an externally accessible, supported lipid membrane, allowing membrane-proximal binding events to regulate reporter assembly without requiring analyte transport across a bilayer.

Additionally, a practical biosensor benefits from a molecular mechanism that produces a specific, low-background signal. Chemically induced dimerization (CID) can couple ligand recognition to protein association^13^. When the interacting partners are fused to split fluorescent reporters with low spontaneous complementation, ligand-induced association can generate fluorescence with low background in the absence of the inducing ligand^14,15^. To this end, we displayed CID-based sensing domains on silica bead-supported lipid bilayers, known as SUPER templates^16^. These structures preserve a membrane-like interface for organizing reporter components while providing a physically supported surface that can be directly exposed to binding partners, washed, and analyzed by fluorescence microscopy.

The FKBP–FRB pair provides a well-characterized module for rapamycin-induced protein association^13^ and has been incorporated into synthetic membrane receptors that couple ligand-induced receptor association to gene expression in mammalian cells.^17^ Here, using membrane-localized FRB/FKBP-mediated split sfCherry reconstitution and antibody-dependent attenuation of fluorescence, we establish a surface-localized model reporter that generates fluorescence in the presence of a dimerizing ligand and reduced fluorescence when an inhibitory binding partner is present.

First, we evaluated rapamycin and its synthetic analog (rapalog)-dependent split sfCherry reconstitution mediated by FRB and FKBP. To do this, we fused FRB and FKBP to complementary fragments of a split sfCherry protein (containing β-strands 1–10 and β-strand 11, respectively)^18,19^ separated by flexible linkers. These split fragments are non-fluorescent individually and exhibit relatively weak effective association. Therefore, interactions that bring them into proximity can enhance fluorescent protein fragment complementation^18,19^.

We co-expressed FRB-sfCherry_1-10_ and FKBP-sfCherry_11_ using a HeLa lysate-based CFE system and evaluated reporter reconstitution under three conditions (**Fig. 1a**): the addition of rapalog, rapamycin, or water. We observed sfCherry reconstitution, indicated by a strong red fluorescent signal, in the rapalog-treated condition. This signal was notably higher than in the rapamycin-treated condition, while the water control yielded almost no fluorescence (**Fig. 1b**). Due to its superior performance in promoting reporter reconstitution in this context, we selected rapalog for all subsequent experiments.

**Fig 1.**
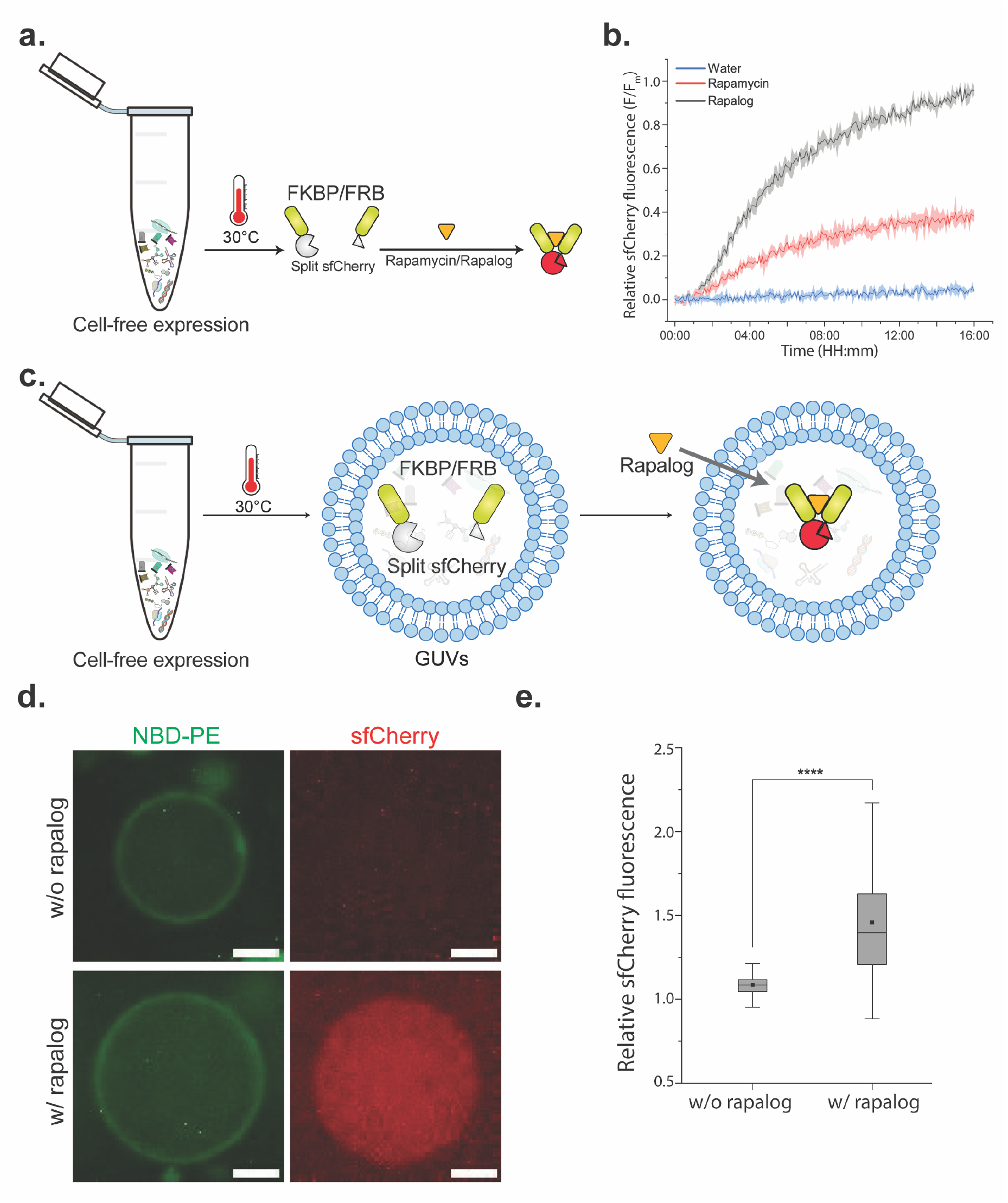
Rapalog-dependent FRB/FKBP association drives functional split sfCherry reconstitution in a mammalian CFE system. (a) Schematic of split sfCherry reconstitution promoted by rapalog- or rapamycin-induced FRB/FKBP association. (b) Relative fluorescence kinetics after addition of 150 nM of rapalog (gray), rapamycin (red), or water (blue). Values represent the ratio of the sfCherry signal to the maximum signal recorded in each independent experiment. Data are shown as mean ± SD (n = 3). (c) Schematic showing bulk CFE of FRB-sfCherry_1–10_ and FKBP-sfCherry_11_, subsequent encapsulation of the resulting reaction mixture in GUVs, and external addition of rapalog. (d) Representative confocal images showing GUVs with or without rapalog (150 nM) treatment. NBD-PE marks the membrane, and sfCherry indicates reporter reconstitution. Scale bars: 5 µm. (e) Quantification of relative luminal sfCherry fluorescence in GUVs. Values represent the ratio of luminal sfCherry signal to background signal. n = 432 GUVs were analyzed in total from three different biological replicates. Boxes represent the 25th to 75th percentiles, with the central line indicating the median, and the black square showing the average (**** represents *p* < 0.0001).

As a step towards constructing cell-like reporter systems, we first co-expressed FRB-sfCherry_1-10_ and FKBP-sfCherry_11_ in bulk using the CFE system and then encapsulated the resulting reaction mixture within GUVs. Membrane-permeable rapalog^20^ was subsequently added to the surrounding solution to enable diffusion into the GUVs lumens (**Fig. 1c**). Confocal microscopy and analysis showed a robust sfCherry signal inside GUVs after rapalog addition (**Fig. 1d-e**).

Having successfully reconstituted the split sfCherry system, we hypothesized that introducing a bulky binder adjacent to the dimerization interface could attenuate reporter assembly and, consequently, reduce the fluorescent signal through steric interference. As a proof of concept, we used the N-terminal 3xFLAG tags on both reporter constructs (3xFLAG-FRB-sfCherry_1-10_ and 3xFLAG-FKBP-sfCherry_11_) as binding sites for anti-FLAG antibodies. We co-expressed these constructs in the CFE system and quantified the resulting sfCherry fluorescence using a plate reader across varying concentrations of both rapalog and an anti-FLAG antibody (**Fig. 2a**). Consistent with our hypothesis, the sfCherry signal decreased in a dose-dependent manner with increasing anti-FLAG antibody concentrations or decreasing rapalog concentrations, indicating that antibody preincubation before rapalog addition can attenuate dimerization-induced reporter output (**Fig. 2b**). Importantly, constructs lacking the 3xFLAG tag retained substantial rapalog-induced fluorescence in the presence of anti-FLAG antibody **(Fig. S1)**. This control is consistent with a contribution from 3xFLAG/anti-FLAG binding to reporter attenuation. Additionally, to show that this system is not limited to anti-FLAG, we replaced 3xFLAG with 5xHis, generating 5xHis-FKBP-sfCherry_11_ and 5xHis-FRB-sfCherry_1-10_. By co-expressing 5xHis-FKBP-sfCherry_11_ and 5xHis-FRB-sfCherry_1-10_, we confirmed sfCherry reconstitution in the presence of rapalog (**Fig. S2**), and fluorescence was reduced in the presence of a Penta-His antibody (**Fig. S3**). Together, these results are consistent with a model in which antibody binding near the FRB/FKBP split sfCherry interface attenuates productive reporter reconstitution through steric interference.

**Fig 2.**
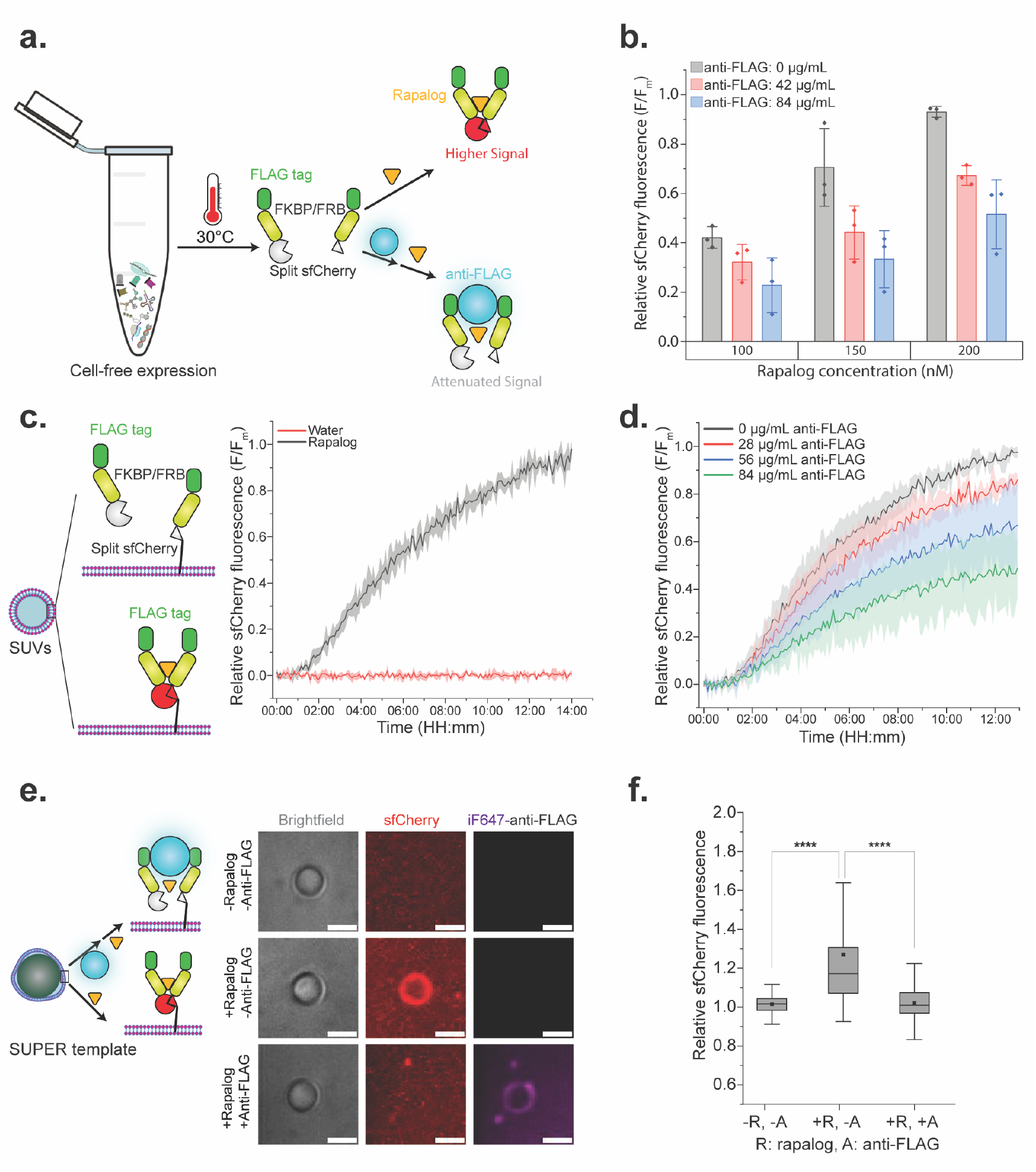
Anti-FLAG antibodies reduce the reconstituted split sfCherry signal by binding to the adjacent 3xFLAG tag and attenuating FKBP/FRB-mediated reporter reconstitution. (a) Schematic of antibody-mediated steric inhibition of 3xFLAG-FRB/FKBP-driven split sfCherry reconstitution. (b) Bulk CFE relative fluorescence response across rapalog and anti-FLAG antibody concentrations. Values represent the ratio of the sfCherry signal to the maximum signal recorded in each independent experiment. Data represent mean ± SD (n = 3), with individual values plotted as points. (c) Left: Schematic of SUV-based assay showing rapalog-dependent recruitment of soluble 3xFLAG-FRB-sfCherry_1-10_ to membrane-anchored 3xFLAG-FKBP-sfCherry_11_-TfR. Right: relative fluorescence of reconstituted sfCherry in SUV-containing reactions following the addition of 150 nM rapalog (gray) or water (red). Values represent the ratio of the sfCherry signal to the maximum signal recorded in each independent experiment. Data are presented as mean ± SD (n = 3). (d) Anti-FLAG antibody dose-dependent attenuation of sfCherry reconstitution in SUV-containing reactions. Values represent the ratio of the sfCherry signal to the maximum signal recorded in each independent experiment. Data are presented as mean ± SD (n = 3). (e) Representative confocal images of SUPER templates under the indicated rapalog (150 nM) and anti-FLAG antibody (84 μg/mL) conditions. sfCherry is shown in red and iF647-anti-FLAG in magenta. Scale bars: 5 µm. (f) Quantification of membrane-localized sfCherry fluorescence on SUPER templates. Values represent the ratio of membrane sfCherry signal to background signal. n = 735 SUPER templates were analyzed in total from three different biological replicates. Boxes represent the 25th to 75th percentiles, with the central line indicating the median, and the black square showing the average (**** represents *p* < 0.0001).

As detecting a non-membrane-permeable ligand requires the sensing domains to be exposed on the outer leaflet of a membrane, we first engineered a membrane-anchored construct by fusing the C-terminus of 3xFLAG-FKBP-sfCherry_11_ to the transmembrane domain of the transferrin receptor (TfR), thus allowing for membrane localization of the 3xFLAG-FKBP-sfCherry_11_ construct^19^.

We initially co-expressed the membrane-anchored 3xFLAG-FKBP-sfCherry_11_-TfR construct with soluble 3xFLAG-FRB-sfCherry_1–10_ in the presence of 100 nm DOPC small unilamellar vesicles (SUVs). We selected SUVs for these validation experiments because they are osmotically robust, can be added directly to CFE reactions to support membrane protein expression, and are compatible with bulk fluorescence measurements using a plate reader. Using a plate reader, we observed that rapalog increased sfCherry fluorescence in reactions containing SUVs and the membrane-anchored construct, consistent with the proposed recruitment mechanism (**Fig. 2c**). Consistent with the bulk CFE reaction, the addition of increasing concentrations of anti-FLAG antibodies resulted in a dose-dependent reduction of the sfCherry signal (**Fig. 2d**).

As a step towards a more practical vesicle-inspired reporter platform, we transferred the membrane-localized reporter from SUVs to SUPER templates. The silica bead support in SUPER templates provides a membrane-coated particle format that can be collected by low-speed centrifugation and washed before analysis ^16^. These handling advantages make bead-supported membranes useful for developing membrane-associated reporter assays. Combined with their uniform 5 µm diameter, these features make SUPER templates attractive candidates for future membrane-associated sensing platforms.

We incubated the SUPER templates in the CFE system containing plasmids for both the membrane-anchored and soluble constructs, and then tested them under three conditions: 1) in the presence of both rapalog and anti-FLAG antibodies, 2) in the presence of rapalog without anti-FLAG antibodies, and 3) a control lacking both rapalog and anti-FLAG antibodies.

sfCherry signal localized to the SUPER template membrane was observed in the rapalog-only condition, as shown in **Fig. 2e-f**, consistent with anti-FLAG-mediated attenuation of the fluorescence output through steric interference. Anti-FLAG antibody recruitment to membrane-localized 3xFLAG-tagged reporter components was confirmed by iFluor 647 fluorescence (i.e., anti-FLAG antibody is labeled with iFluor 647).

To advance towards a fully membrane-localized reporter design, we appended a transmembrane domain to the second fragment, constructing 3xFLAG-FRB-sfCherry_1-10_-TfR. After confirming that both membrane-anchored fragments supported rapalog-dependent sfCherry reconstitution in SUVs and showed dose-dependent anti-FLAG-mediated attenuation of fluorescence (**Fig. 3a-b**), we evaluated this fully membrane-anchored system on the SUPER template platform. As shown in **Fig. 3c-e**, membrane-localized sfCherry fluorescence was observed robustly when rapalog was added in the absence of anti-FLAG antibodies. This supports the functional membrane localization of both 3xFLAG-FRB-sfCherry_1-10_-TfR and 3xFLAG-FKBP-sfCherry_11_-TfR, as well as antibody-dependent attenuation of rapalog-induced reporter fluorescence, thereby establishing a model reporter that produces a high signal when the dimerizing ligand is present and a reduced signal when the inhibitory binding partner is present.

**Fig 3.**
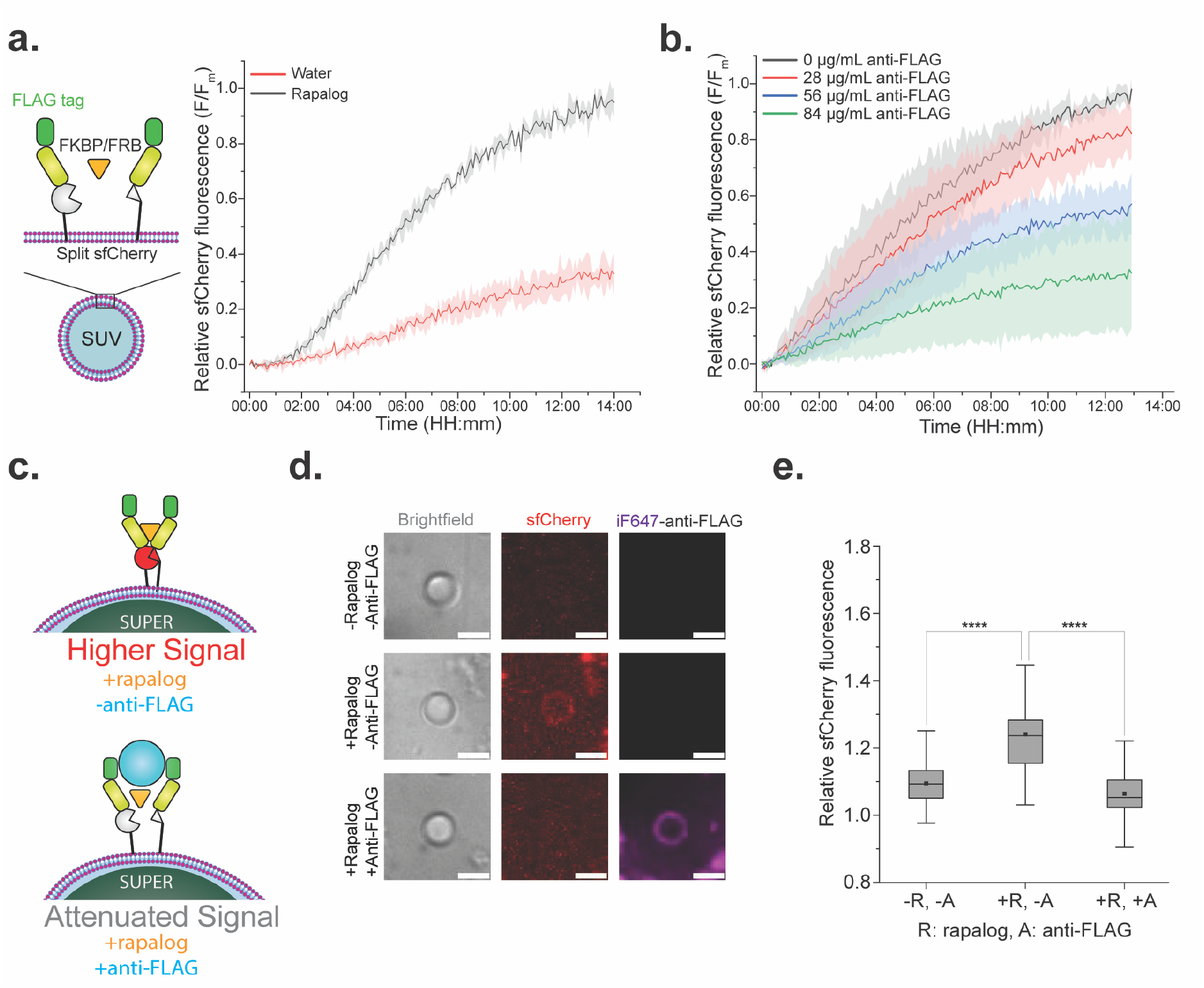
SUPER templates support antibody-modulated split sfCherry reconstitution using membrane-anchored 3xFLAG-FKBP/FRB constructs. (a) SUV assay showing rapalog-dependent (150 nM) reconstitution of two membrane-anchored split sfCherry constructs. Values represent the ratio of the sfCherry signal to the maximum signal recorded in each independent experiment. Data are presented as mean ± SD (n = 3). (b) Anti-FLAG antibody dose-dependent attenuation of sfCherry in SUV-containing reactions. Values show the ratio of the sfCherry signal to the maximum signal recorded in each independent experiment. Data are presented as mean ± SD (n = 3). (c) Schematic of the SUPER-template reporter configuration: rapalog induces sfCherry reconstitution, while anti-FLAG antibody attenuates reporter reconstitution. (d) Representative confocal images of SUPER templates under the indicated rapalog (150 nM) and anti-FLAG antibody (84 μg/mL) conditions. sfCherry is shown in red and iF647-anti-FLAG in magenta. Scale bars: 5 µm. (e) Quantification of membrane-localized sfCherry fluorescence on SUPER templates. Values represent the ratio of membrane sfCherry signal to background signal. n = 294 SUPER templates were analyzed in total from three different biological replicates. Boxes represent the 25th to 75th percentiles, with the central line indicating the median, and the black square showing the average (**** represents *p* < 0.0001).

While the current study uses the 3xFLAG tag and anti-FLAG antibody as a model recognition pair to demonstrate antibody-dependent attenuation of reporter output, the platform could in principle be adapted to other binding pairs for future detection of applicable targets. We anticipate that sufficiently bulky molecules that bind near the dimerization interface will be able to attenuate fluorescence output, although the efficiency of signal attenuation will depend on binding affinity, epitope position, linker length, and target size. By substituting the 3xFLAG or 5xHis tag with specific epitopes, receptor domains, or aptamers, this architecture could provide a basis for future studies toward sensors targeting viral antigens, cancer-associated circulating proteins, antibodies, or environmental toxins. Importantly, these can all be coupled to the well-established FKBP/FRP dimerization domains, making the steric attenuation of membrane-associated fluorescence a universal detection platform simply by swapping out the epitope tags. Furthermore, because the sensing domains are localized to the outer leaflet of the structurally robust SUPER template, this architecture may be useful for future detection of membrane-impermeable analytes without requiring target transport across a lipid bilayer.

In complex applications, relying on a single binding event often lacks the specificity required for accurate detection. By engineering an architecture based on the combination of ligand-induced complementation and a signal-attenuating binding partner, our platform introduces a programmable layer of regulation. Future studies could establish target specificity and distinguish target-dependent attenuation from nonspecific signal loss.

In summary, these results demonstrate antibody-dependent attenuation of dimerization-induced reporter fluorescence in cell-free membrane systems, consistent with a steric interference mechanism, providing a proof-of-concept platform for future molecular engineering and biosensing applications. The supported membrane format may improve structural robustness relative to free-standing vesicles and could facilitate washing and handling steps prior to signal readout.

## Acknowledgement

We thank Sung-Won Hwang for her help with the experiments and valuable insights. We also thank Samuel Chen, other members of the Liu Lab, and Cheemeng Tan (University of California, Davis) for helpful discussions. A.P.L. acknowledges support from the National Institutes of Health (R21 AI173559 and R01 GM144652). A.P.L., P.P., and H.M. conceptualized the study. P.P., H.M., and A.P.L. designed the experiments. P.P. and H.M. conducted the experiments. P.P. performed image processing. All authors contributed to drafting the manuscript and reviewed and approved the final version.

## Data availability

The data supporting this study, including detailed materials and methods, additional figures, and amino acid sequences of the protein components, are included in the Supplementary Information.

