## Supplementary Information for "Steric attenuation of membrane-associated dimerization-induced fluorescence for cell-free biosensing"

**Materials and Methods:**

Cloning and DNA construct preparation

Plasmids encoding FKBP and FRB were a gift from Aaron Morris (University of Michigan). Plasmids encoding the transferrin receptor transmembrane domain, sfCherry$_{1\text{-}10}$, and sfCherry$_{11}$ were from our previous study[1]. The target gene sequences were amplified from template plasmids via PCR using Q5 High-Fidelity 2X Master Mix (NEB, M0492S). All primers were purchased from Eurofins Scientific. Amino acid sequences of protein components used in this study are summarized in **Table S1**.

For cell-free expression (CFE), the pT7CFE1-6xHis vector (Thermo Fisher Scientific) was used as the backbone. To preserve the hydrophobicity of the C-terminal regions of the expressed proteins, the 6xHis tag was removed from the vector for all constructs.

Target plasmids were constructed from gene fragments using Gibson assembly, and the final DNA constructs were verified using either Sanger or Nanopore sequencing (Eurofins Genomics). The assembled DNA constructs were purified from DH5α *E. coli* (NEB) using a miniprep kit (Qiagen).

We first generated 3xFLAG-FKBP-sfCherry$_{11}$ and 3xFLAG-FRB-sfCherry$_{1–10}$ to assess rapalog-induced sfCherry reconstitution in bulk CFE reactions and GUVs (**Fig. 1**). For membrane-associated assays, we added a TfR-derived membrane anchor to each construct, generating 3xFLAG-FKBP-sfCherry$_{11}$-TfR and 3xFLAG-FRB-sfCherry$_{1–10}$-TfR. Constructs lacking the 3xFLAG tag, FKBP-sfCherry$_{11}$ and FRB-sfCherry$_{1–10}$, were prepared as controls for nonspecific effects of anti-FLAG antibody. Finally, we replaced 3xFLAG with 5xHis to generate 5xHis-FKBP-sfCherry$_{11}$ and 5xHis-FRB-sfCherry$_{1–10}$ for experiments with Penta-His antibody.

SUV preparation

SUVs were generated via thin-film hydration and extrusion. 78.6 µL of a 25 mg/mL solution of 1,2-dioleoyl-sn-glycero-3-phosphocholine (DOPC; Avanti Polar Lipids) in chloroform was dried under a gentle argon stream and desiccated at room temperature for 1 h to remove residual chloroform. The resulting lipid film was hydrated using 500 µL of ultra-pure water and vortexed to achieve complete resuspension. SUVs were prepared by extruding the suspension 11 times through a 100-nm filter using an Avanti mini-extruder. The final solution was stored at 4 °C and utilized within a week.

SUPER template preparation

SUPER (supported bilayers with excess membrane reservoir) templates were prepared according to a previously published protocol[2] with slight modifications. 50 µL of the SUV dispersion and 4 µL of 5 µm silica beads (Bangs Laboratories; ~2.85 × $10^6$ beads) were combined in a 100 µL reaction mixture containing 1 M NaCl. The suspension was incubated at room temperature for 30 min with occasional gentle flicking. After incubation, the solution was diluted to 1 mL using PBS (Thermo Fisher), and the beads were collected via centrifugation at

200 × g for 2 min. Following an additional PBS wash under the same conditions, the final SUPER templates were resuspended in 30 µL of ultra-pure water.

GUV preparation

GUVs were prepared via an inverted emulsion protocol[3] with a few modifications. Chloroform-dissolved lipid stocks of 1-palmitoyl-2-oleoyl-glycero-3-phosphocholine (POPC), cholesterol, and 1,2-dioleoyl-sn-glycero-3-phosphoethanolamine-N-(7-nitro-2-1,3-benzoxadiazol-4-yl) (NBD-PE) (Avanti Polar Lipids) were mixed in a glass vial at a 79.8:20:0.2 molar ratio. Following chloroform evaporation under a gentle argon stream, mineral oil (Sigma-Aldrich, cat#: M5904) was added to yield a 500 µM final lipid concentration. Lipid dissolution in oil was ensured by thorough vortexing, followed by incubation at 55 °C for 20 min, and vortexing a second time. Next, an oil-water interface was made in a 1.5 mL microcentrifuge tube by layering 300 µL of the lipid-in-oil solution over 400 µL of the outer solution. An outer-buffer stock solution containing 1.7 M HEPES-KOH and 0.5 M Tris was diluted with water to match the measured osmolality of the inner solution. After a 1-hour room-temperature incubation of the interface, a uniform emulsion was made in a separate tube by vigorously pipetting 600 µL of the lipid-in-oil solution with 11 µL of an inner solution. This inner solution was composed of a 10 µL post-expression CFE reaction and 1 µL OptiPrep (Sigma-Aldrich, cat#: D1556). Here, OptiPrep serves to create a density gradient (the osmolality of both the inner and outer solutions was measured using an osmometer, and the outer solution was diluted with water to achieve osmotic balance). The emulsion was transferred to the top of the oil-water interface and centrifuged at 2800 × g for 10 min at room temperature to drive GUV formation. Finally, the top 900 µL of the oil phase and the bulk of the outer solution were aspirated until approximately 50 µL of outer solution remained, after which the pelleted GUVs were resuspended by gentle pipetting.

CFE reactions

Proteins were synthesized using the 1-Step Human Coupled In Vitro Transcription-Translation Kit (Thermo Fisher, cat#: 88881). Briefly, 5 µL of HeLa lysate was combined with 2 µL of reaction mix and 1 µL of accessory proteins. The final reaction volume was brought to 10 µL with the addition of 0.5 µL of RNase inhibitor (NEB, cat# M0314S), 400 ng of plasmid DNA (200 ng per plasmid), and ultra-pure water. For membrane-based sfCherry reconstitution assays, the water fraction was replaced with the appropriate SUV or SUPER template suspension. Bulk or SUV-containing CFE reactions were incubated at 30 °C for 5 h. CFE reactions containing SUPER templates from **Fig. 2e-f** and **Fig. 3c-e** were incubated for 8 and 3.5 hours, respectively.

GUV and bulk CFE assays

Rapalog (A/C heterodimerizer; TaKaRa, cat#: 635057), rapamycin (BPS Bioscience, cat#: 27062), or ultra-pure water was added to the GUV suspension or bulk CFE reaction (**Fig. 1a-b**). Following an 8 h incubation at room temperature, the GUVs were transferred to Corning clear glass-bottom 96-well plates for imaging.

Antibody treatments and sfCherry reconstitution assays

In experiments utilizing anti-FLAG antibodies (DYKDDDDK tag antibody-iFluor647; GenScript, cat#: A01811) or Penta-His antibodies (Penta·His Alexa Fluor 647 Conjugate; Qiagen, cat#: 35370), the antibody was added directly to the completed CFE reaction, with controls receiving an equivalent volume of PBS. At 30 minutes post-antibody addition, rapalog was added; then, reactions were transferred to Corning low-volume 384-well plates, and bulk sfCherry fluorescence was measured on a Cytation 5 plate reader (BioTek) using 579 nm excitation and 616 nm emission wavelengths. For confocal microscopy of the SUPER templates, the samples were incubated for an additional 8 h or 4 h (**Fig. 2e-f** and **Fig. 3c-e**, respectively) following the addition of rapalog at room temperature. The SUPER templates were then diluted 1:5 with PBS (Thermo Fisher) and transferred to Corning low-volume clear glass-bottom 384-well plates for imaging.

Plate reader assays

In all plate reader assays, the first five fluorescence measurements (0–20 min) were averaged for each condition. This baseline was then subtracted from every measurement in the corresponding condition. Fluorescence was normalized to the maximum signal recorded among all the different conditions for each biological replicate.

Confocal imaging and image analysis

Fluorescence images were acquired on an Olympus IX-81 inverted microscope equipped with a Yokogawa CSU-X1 spinning disk confocal unit, an iXON3 EMCCD camera (Andor Technology), a National Instruments DAQ-MX-controlled laser unit (Solamere Technology), and a 60x/1.4 NA oil-immersion Plan-Apochromat objective (Olympus). All image acquisitions were managed using MetaMorph software (Molecular Devices). NBD, sfCherry, and iFluor647 fluorescence were excited at 488 nm, 561 nm, and 640 nm, respectively.

Images were analyzed using custom Python code developed with assistance from OpenAI Codex (GPT-6 Astra), with manual review and adjustments, followed by manual cross-checking in ImageJ. For SUPER templates, beads were identified in DIC images and manually reviewed. Mean membrane fluorescence (M) was measured within an annulus with inner and outer diameters of 17 and 25 pixels, respectively (4.3 pixels = 1 μm), including only pixels closer to the target bead than to any selected neighboring bead. Local background fluorescence (B) was measured within a concentric annulus with inner and outer radii of 18 and 23 pixels, excluding pixels within 16 pixels of selected neighboring bead centers. Relative sfCherry fluorescence was calculated as (M−100)/(B−100), where 100 analog-to-digital units (ADU) was the camera offset. For GUVs, candidates were identified from the 488-nm membrane-dye channel and manually reviewed, including candidates with faint visible membranes. Candidates were retained if their fitted area-equivalent diameter was >5 μm and their minor-to-major axis ratio was >0.90. Mean luminal reporter intensity (L) was measured from the original 561-nm image within a circular ROI whose radius equaled the fitted minor semiaxis minus 3 pixels (4.3 pixels = 1 μm). Mean local background intensity (B) was measured within the nearest eligible circular ROI of 8-pixel radius, avoiding detected vesicles and other structures. Relative luminal sfCherry fluorescence was calculated as (L−100)/(B−100), where 100 ADU was the camera offset. Quantitative image data were exported to Microsoft Excel. Statistical significance was determined using a two-tailed t-test assuming unequal variances to calculate *p*-values with Excel's built-in data analysis tool.

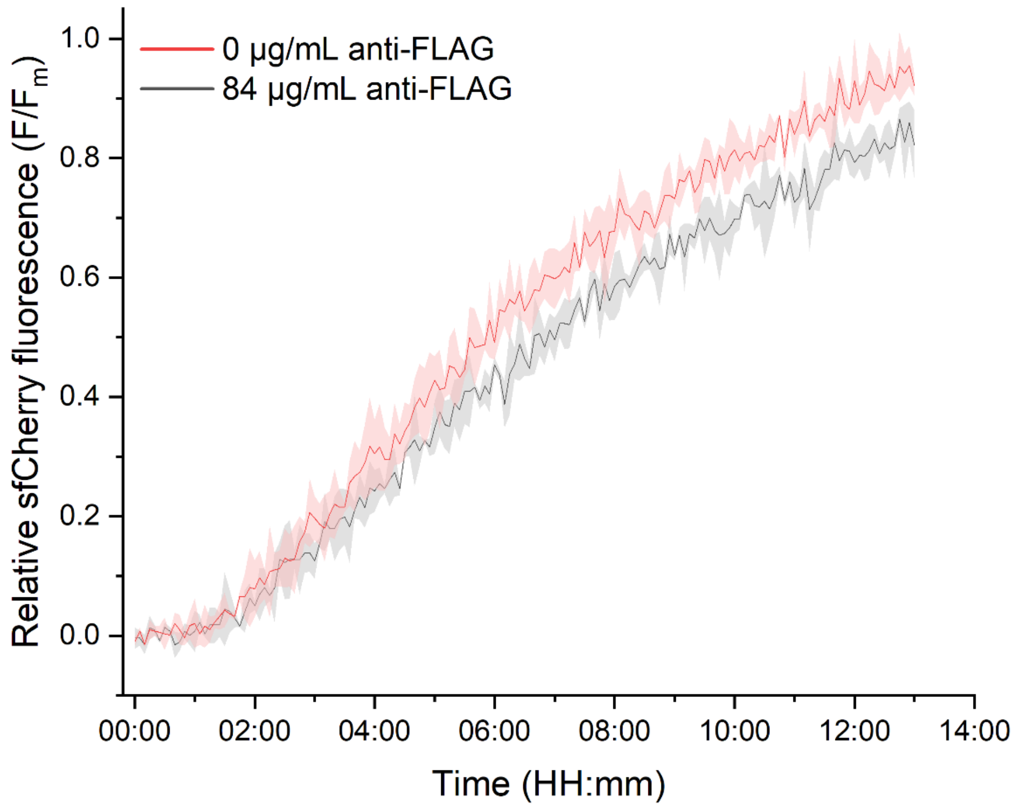


**Fig. S1. Effect of anti-FLAG antibody on split-sfCherry reconstitution in untagged constructs.** Relative sfCherry fluorescence in bulk cell-free reactions co-expressing FRB-sfCherry$_{1–10}$ and FKBP-sfCherry$_{11}$, with 84 µg/mL anti-FLAG antibody (gray) or without antibody (red), following the addition of 150 nM rapalog. Fluorescence was normalized to the maximum signal recorded in each independent experiment. Data are presented as mean ± SD (n = 3).

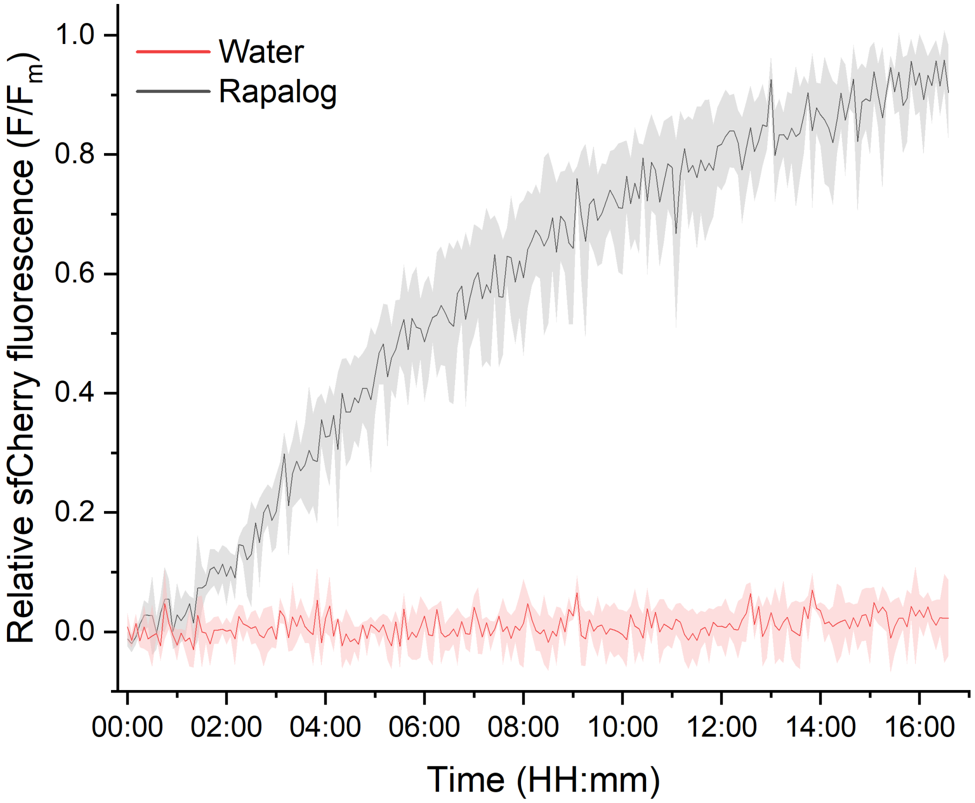


**Fig. S2. Rapalog promotes split-sfCherry reconstitution in 5xHis-tagged constructs.** Relative fluorescence of reconstituted sfCherry in bulk of co-expressed 5xHis-FKBP-sfCherry$_{11}$ and 5xHis-FRB-sfCherry$_{1-10}$ following the addition of 150 nM rapalog (gray) or water (red). Fluorescence was normalized to the maximum signal recorded in each independent experiment. Data are presented as mean ± SD (n = 3).

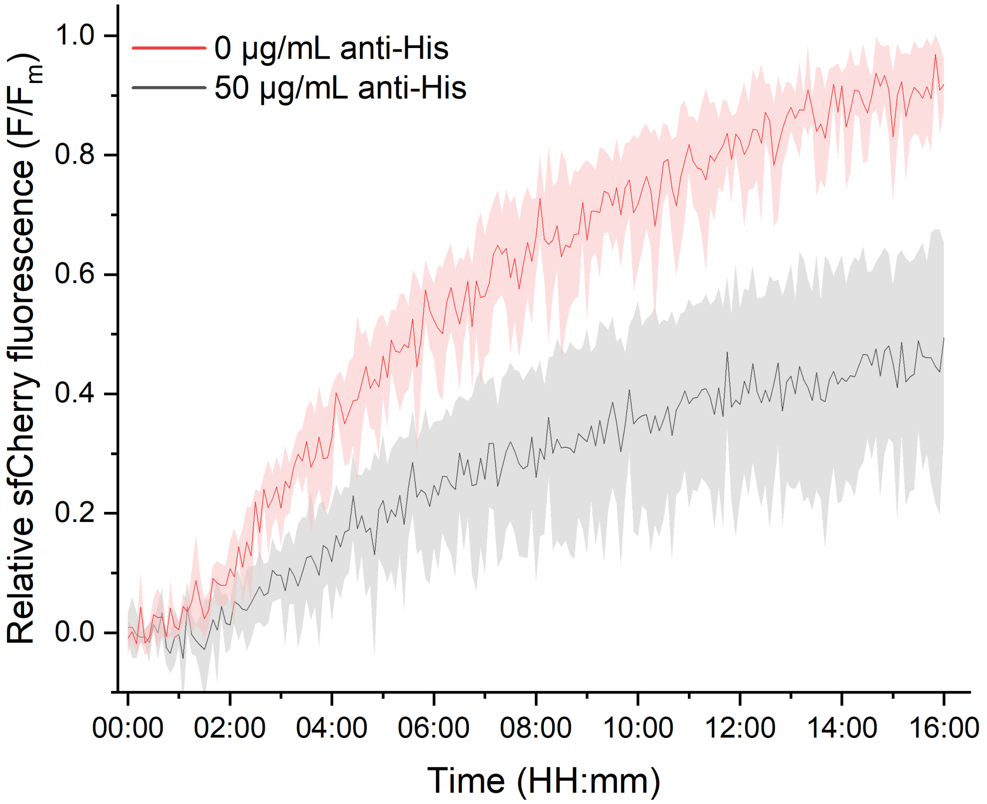


**Fig. S3. Anti-His antibody attenuates fluorescence in 5xHis-tagged split-sfCherry constructs.** Relative sfCherry fluorescence in bulk cell-free reactions co-expressing 5xHis-FRB-sfCherry$_{1–10}$ and 5xHis-FKBP-sfCherry$_{11}$, with addition of 50 µg/mL Penta-His antibodies (gray) or without antibody (red), following the addition of 150 nM rapalog. Fluorescence was normalized to the maximum signal recorded in each independent experiment. Data are presented as mean ± SD (n = 3).

| Table S1. Amino acid sequences of the protein components and tags used in this study. | |
|---|---|
| FKBP | MGVQVETISPGDGRTFPKRGQTCVVHYTGMLEDGKKFDSSRDRNKPFKFML<br>GKQEVIRGWEEGVAQMSVGQRAKLTISPDYAYGATGHPGIIPPHATLVFDVEL<br>LKLE |
| FRB | MLEMWHEGLEEASRLYFGERNVKGMFEVLEPLHAMMERGPQTLKETSFNQ<br>AYGRDLMEAQEWCRKYMKSGNVKDLLQAWDLYYHVFRRISKTR |
| $sfCherry_{1-10}$ | MEEDNMAIIKEFMRFKVHMEGSVNGHEFEIEGEGEGHPYEGTQTAKLKVTKG<br>GPLPFAWDILSPQFMYGSKAYVKHPADIPDYLKLSFPEGFTWERVMNFEDGG<br>VVTVTQDSSLQDGQFIYKVKLLGINFPSDGPVMQKKTMGWEASTERMYPED<br>GALKGEINQRLKLKDGGHYDAEVKTTYKAKKPVQLPGAYNVDIKLDITSHNED |
| $sfCherry_{11}$ | YTIVEQYERAEARHST |
| TfR | MDQARSAFSNLFGGEPLSCTRASLARQVDGDNSHVEMKLAVDEEENADNNT<br>KANVTKPKRCSGSICYGTIAVIVFFLIGFMIGYLGY |
| 3xFLAG | DYKDHDGDYKDHDIDYKDDDDK |
| 5xHis | HHHHH |
